# Byte-exact, format-aware compression for FASTA, FASTQ, BED and VCF files

**DOI:** 10.64898/2026.03.16.712193

**Authors:** Michail Patsakis, Alexandros Margaris, Theodore Chronopoulos, Ioannis Mouratidis, Ilias Georgakopoulos-Soares

## Abstract

**Motivation:** Most omics data is stored by compressors that treat it as an unstructured byte stream. Format-aware compressors close that gap but are fragmented across formats and often expect a training pass over the user’s own file. A general-purpose lossless compressor that is applicable to the most storage-intense omics data formats while beating the general codecs on ratio and speed would offer significant savings for large omics datasets storage and data retrieval.

**Results:** We present NYX, a compressor for FASTA, FASTQ, BED and VCF files. It leverages a reversible, format-specific transform optimised for the OpenZL compression algorithm to reach compression ratios up to 2.3 times those of the strongest general-purpose codecs (zstd -19, xz -9e and 7-Zip) while compressing up to two orders of magnitude faster. On the 1000 Genomes VCF dataset NYX reaches a compression ratio of 468.67x, 2.0 times that of xz -9e while compressing 1.8 times faster. On mammalian FASTA assemblies totalling 9.3 GB, NYX is within 3-5% of NAF, the strongest purpose-built FASTA codec, while compressing 57-73 times faster. Across all 54 inputs no single codec beats NYX on ratio and compression speed together: the tools that achieve higher compression are 1.2 to 73 times slower. NYX provides fully lossless, fast and efficient compression under one easy-to-use tool for the most common omics data types.

**Availability and implementation:** Source code, the shipped models, the benchmark harness and the scripts that regenerate every figure and table: https://github.com/Georgakopoulos-Soares-lab/compression

## 1 Introduction

Public archives such as the Sequence Read Archive, the European Nucleotide Archive and the China National GeneBank now collectively host tens of petabytes of raw and processed data [1, 2], and continue to grow: the ENA alone doubles its sequenced nucleotides roughly every 45 months [3]. Large sequencing centers routinely generate tens of terabytes per day, cumulatively producing tens of petabytes per year. This unprecedented output has transformed genomics into one of the fastest-growing domains of the global datasphere, placing substantial demands on storage, transfer, and computation infrastructures [4]. At this scale, storage and transfer costs directly constrain data sharing, reproducibility, and secondary analysis.

To manage this data deluge, a stable ecosystem of file formats has emerged, each matched to a stage of the analysis. FASTA and FASTQ store raw reads and sequence metadata [5, 6], SAM, BAM and CRAM encode alignments with increasingly space-efficient binary representations [7, 8], VCF and its binary form BCF carry variant calls, BED carries interval annotations, and BigWig, BigBed and HDF5-based schemas hold quantitative signals such as coverage, expression and methylation. These formats are highly structured: constrained alphabets, fixed field templates, and columnar redundancy across records are properties of the specifications themselves, and are therefore available to any compressor that parses them.

Because this structure is a property of the specifications, a compressor that parses the format can exploit it. Genozip [9, 10], SPRING [11] and NAF [12] achieve substantially higher ratios than general-purpose codecs on the formats they target, and the size of that gap is a direct measure of how much redundancy the formats expose. The gain comes at a cost, however. At their strongest settings these tools are often one to two orders of magnitude slower than general-purpose codecs. The open-source tools each serve a single format or a small group of them, so a group working across raw reads, assemblies and variant calls must install, learn and maintain several unrelated tools; Genozip spans the formats but is closed-source and commercially licensed, placing it beyond the reach of many laboratories. Faced with that trade, most laboratories and archives default to gzip [13], which is fast, free and universal but treats every file as an unstructured byte stream: it cannot separate records into independently modelled sub-streams, and therefore forfeits the redundancy that the formats make available. No existing tool combines this redundancy gain with speed, open availability and coverage of all four formats, which is the gap NYX is designed to fill.

A framework that is to serve all of these formats has to satisfy two primary requirements:

First, a learned model is sensitive to the gap between its training data and its target. A configuration trained on a file structurally similar to the target performs best, but that presumes the user can supply representative training data and absorb a per-file optimisation cost.

Second, performance varies across subtypes within a nominal format. VCF is the clearest case. The specification admits sites-only annotation sets, single-sample germline calls, gVCFs dominated by reference blocks, tumour–normal somatic sets, family pedigrees, joint-genotyped cohorts and panels with thousands of columns. These share a grammar but differ in where their redundancy lies, and a model tuned for one fits another poorly.

NYX addresses both. Compression is performed against a fixed set of pre-trained OpenZL graphs, and no training is performed on user data. The model training algorithm is included for users wishing to optimize for their specific data, but is not required to achieve state-of-the-art performance on the majority of common datasets. For VCF the dominant cost is the genotype matrix, which is most of the bytes in any cohort or panel file and almost none of them in a single-sample call set. Rather than dispatch between models per file subtype, NYX applies one transform whose components each pay off where they apply: a haplotype-level positional Burrows-Wheeler transform on the genotype matrix, per-INFO-key and per-FORMAT-subfield streams, and a set of value rewrites for integers, coordinates, fixed-point decimals, integer lists and low-cardinality text (section 3.3). Models are trained per stream class rather than per file type, so a file carrying INFO or FORMAT keys absent from the training corpus still routes every stream to a shipped model. NYX also lets the user cap peak memory directly so a 28 GB file compresses within the same footprint as a 1 GB one.

Every figure we report is paired with a full decompression and a byte-for-byte comparison against the original. Section 3 reports compression ratio, throughput and peak memory for FASTA, FASTQ and VCF against general-purpose codecs and, where available, format-specific tools, and Section 3.5 examines the trade-off between ratio and speed across all inputs.

## 2 Methods

### 2.1 Core Pipeline & OpenZL Substrate

Each format follows a three-stage pipeline: a reversible, format-specific transform rewrites the file into homogeneous streams. A fixed OpenZL [14] graph compresses those streams, and the output is chunked for parallel, independent decoding. We use a custom version of OpenZL 0.2.5 which raises the encoder input and stream-dispatch limits from 2048 to 4096.

Training is an offline, maintainer-side step. No sampling or model fitting is performed on user data. Because plans are sensitive to the distance between training and target data, shipped models are trained on heterogeneous corpora. If a structural mismatch occurs, pipelines fall back to a generic OpenZL profile. The training corpora are disjoint from the benchmark corpus of this paper: the FASTA models are trained on six assemblies (E. coli, yeast, Arabidopsis, macaque, rat and barley), none of them benchmarked, and the FASTQ models on the first 40,000 reads of each of eight public sequencing runs, none from a study that contributes a benchmark file. The BED codec ships no model at all. The one overlap is in VCF, where the CIViC annotation file is both a training input and a benchmark file; the shipped models change that file’s ratio by 0.1% (Section 3.3.3), so the overlap cannot materially affect its result. The accessions and the training scripts are in the repository.

### 2.2 Format-Specific Transforms

#### 2.2.1 FASTA Transform

A FASTA record conflates distinct data types that compress efficiently in isolation but poorly together. The NYX container explicitly separates these components into independent streams to maximize compression:

- Nucleotide sequences are isolated and immediately reduced to 2 bits per base.
- Soft-masking and sequence exceptions (such as non-ACGT characters and IUPAC codes) are extracted as structural run lengths, shrinking megabase-spanning gaps to just a few bytes.
- File formatting and headers, including verbatim text and arbitrary line-wrapping, are preserved independently at near-zero byte cost.

The transform is byte exact by construction. Soft-masking, IUPAC ambiguity codes, arbitrary line widths, CRLF line endings, and a missing trailing newline all round-trip. There is no sequence-only or case-insensitive mode.

Input is cut on record boundaries into chunks of at most 450 MiB; a single record longer than that, such as a wheat chromosome, forms a chunk of its own. The cap is NYX’s and bounds the memory each worker holds; OpenZL itself sets no limit on the size of an input it compresses. Its limits are elsewhere: on the number of streams in one frame (2048, which we raise to 4096 for the 2,504-sample 1000 Genomes panel) and, when training, on the training samples (by default 150 MiB per sample and 300 MiB in total, with larger samples left out rather than truncated). Our training corpora stay below both. Each chunk is compressed twice, once with the shipped FASTA model and once with an untrained schema profile that splits the packed record into its component streams, and the smaller result is kept; in our runs the model is the smaller on the mammalian assemblies and the schema split on the highly repetitive wheat genome. The shipped model is OpenZL’s serial graph with its parameters set by training. Training it on a corpus that shares no genome with the benchmark (E. coli, yeast, A. thaliana, rhesus macaque, rat and barley) produced a byte-identical model to training on the benchmark assemblies themselves, so the FASTA results do not depend on having seen the test data.

#### 2.2.2 FASTQ Transform

Each FASTQ record is split into identifier, sequence and quality segments. Identifiers are tokenised into alphanumeric and delimiter runs and stored column-wise, with all-digit columns coded as integers and the remainder as text; headers whose token structure varies across reads are stored as raw bytes. Quality scores are stored as a single stream, optionally split by the preceding quality value where this reduces size. To further optimize performance, NYX ships three Illumina FASTQ models, which differ in the reads they were trained on (fixed- or variable-length) and in how the sequence stream is coded (a long-window LZ or zstd), plus two for Nanopore reads. One chunk from the middle of the file is compressed with each shipped model, and the smallest result determines the model used for the whole file. This costs roughly one chunk of compression time, but allows optimal performance. For example, on a small-RNA library, a heuristic choice based on read length and k-mer sharing selected a model that was 16% larger and 1.8 times slower than the one chosen by measurement.

As an optional feature, NYX implements read reordering following HARC [15] and SPRING [11]: reads are sorted by canonical minimizer [16] (k = 15) within a block, and a permutation stream restores the original order. However, it is disabled in all reported runs, as when selecting the optimal compressor among those that NYX ships with, reordering no longer improved performance: specifically on the small-RNA library, where it should help most, it reached 6.15x in 259 s against 6.68x in 62 s without.

#### 2.2.3 VCF Transform

A VCF record interleaves three kinds of data with different statistics: a small set of fixed columns describing a locus, a variable set of key=value annotations, and a genotype matrix that in a cohort or panel file is upwards of 95% of the bytes. The body is cut on record boundaries into independently decodable blocks of a fixed byte budget (64 MB by default), and each block is parsed into named streams.

CHROM, ID, REF, ALT, QUAL and FILTER each become one stream of values. POS becomes a zigzag varint delta; a POS whose text is not a canonical integer is diverted verbatim to an exception stream so no normalisation can occur. The ordered pattern of INFO keys, each flagged for whether it carried ’=’, is interned per block, and every key gets its own value stream; reconstruction walks the pattern and draws one value per key, which reproduces duplicate keys, flag keys and empty values exactly. FORMAT strings are interned likewise, and each non-GT subfield becomes one stream written sample-major.

Where the first FORMAT subfield is GT, each call is parsed into per-haplotype allele codes and the block’s genotype matrix is passed through a haplotype-level positional Burrows-Wheeler transform [17]. At every variant the haplotypes are stably counting-sorted by the alleles seen so far, which places haplotypes sharing long local allele patterns adjacently and reduces each row to a handful of long runs; the permuted row is then run-length coded into separate run-length and run-symbol streams. The permutation is recomputed at decode time from the already-decoded alleles, so nothing about it is stored. Phasing separators and ploidy are stored per variant, and any call that is not a clean fixed-ploidy genotype is diverted to an exception stream with its text intact.

Each text stream is then rewritten by whichever of the following applies, recursively: all-integer streams become fixed-width numerics, or zigzag varint self-deltas where those are smaller, which is what collapses the monotone END coordinates of a gVCF; fixed-point decimals split into sign, integer part, fraction length and fraction value; comma-separated integer lists such as AD and PL split into element counts plus one stream per element position; VEP and SnpEff style annotations split into records and then into one stream per field position; and any remaining low-cardinality text becomes a dictionary plus an index stream.

Each stream is compressed as a separate OpenZL frame, using the shipped model for its stream class. Classes are broad categories such as gt.runlen, text, varint and idx. Streams from keys never seen in training are still assigned a class. For each stream, NYX compares the shipped model with OpenZL’s generic profile and keeps whichever gives the smaller output. This check matters because a model trained on one VCF subtype can do worse on another: applying the shipped models to every stream regardless makes a single-sample VCF 5% larger than using the generic profile. The annotation split is handled the same way. It is skipped when values repeat often enough that the dictionary transform already stores them whole. Section 3.3.3 adds these components one at a time, in this order: the column split, the per-INFO-key streams, the value rewrites, the text dictionaries, the annotation split, the haplotype PBWT and, last, the shipped models.

#### 2.2.4 BED Transform

BED [18] is a family of tab-separated layouts rather than one format. Its first six columns -- chromosome, start, end, name, score and strand -- mean the same thing in every dialect; after column six they do not, so column 7 is thickStart in BED12 and signalValue in the narrowPeak files written by MACS2 [19]. NYX therefore makes no assumption about width. Each block’s modal field count defines its columns, and a row with a different count is stored verbatim on its own, so a stray comment or a ragged line costs that line rather than the block’s format awareness.

Every column is encoded in one of six ways -- as text, as non-negative integers, as row-to-row integer deltas, as a dictionary, as fixed-point decimals with a per-value scale, or as comma-separated integer lists -- or by reference to a column on its left: as an exact copy, as a same-row integer difference, or as a lookup table when the left column determines it. END is START plus a feature length; thickStart and thickEnd repeat START and END whenever a feature has no thick region; itemRgb is a function of a chromatin state’s name. References point only leftwards, which keeps the reference graph acyclic and lets the decoder rebuild a block in one pass.

No encoding is chosen from a guess about what a column holds. For each column, every applicable candidate is built for the first 16,384 rows of the block and compressed, the sizes are projected to the whole block, and the smallest is kept; a lookup table, whose size does not grow with the rows, is priced in full. Columns are priced in parallel, so the choice does not depend on the thread count, and the archive is byte-identical at 1 and 16 threads. Two further settings are measured once per file on an 8 MB prefix of its first block: the OpenZL compression level, taken as the cheapest of {default, 12, 16} whose output is within 2% of the smallest, and whether to also compress each block undivided and keep the smaller result, which is enabled only when the undivided form comes within 3% of the column split on that prefix. The level is not fixed because its effect is not monotone: on a narrowPeak file both level 12 and level 19 produced a larger archive than the default. Blocks are sized from a 4000 MB memory budget by default, and every block is decoded in-process and compared with its input before it is written.

#### 2.2.5 Exactness and robustness

Every block is decoded again in-process and compared against its source bytes before the encoder accepts it; a block that does not reproduce exactly is stored verbatim instead. The transform is therefore exact on any input, and the failure mode is a loss of ratio rather than a loss of data. Anything without a usable #CHROM line, including an empty file, a byte-order mark, CR-only line endings, or a file that is not a VCF at all, is stored through the same verbatim path rather than refused, so the tool is total.

We check this on 155 cases, all byte-exact: 136 legal-but-awkward VCFs run under every combination of the transform switches, and 19 hostile inputs. The hostile set is what a compressor is actually handed: ragged column counts, duplicate sample names, non-ASCII INFO text, symbolic and breakend ALTs, a 949 KB single INFO value, a 640 KB REF/ALT pair, 5000 samples, raw binary, and text that is not a VCF. A first implementation of the verbatim fallback round-tripped two of these to the wrong bytes rather than failing; only the byte comparison caught it, which is the argument for making that comparison unconditional rather than a flag.

### 2.3 Memory Management

NYX allows the user to define available system memory (RAM), with --max-mem-mb N, a budget in megabytes. RAM usage is restricted by implementing the three following features:

- **One Shared Compressor:** The trained graph is deserialized once across all worker threads.
- **Streamed Output:** Compressed frames are written to disk by a bounded in-order writer as they complete.
- **Bounded Blocks:** The chunk size is derived from the user-specified budget, as budget / (threads × 4) for FASTQ, the measured number of copies of a chunk a worker holds. Defaults differ by format because the formats differ in how much memory a chunk costs: FASTA and BED default to a 4000 MB budget, FASTQ to 500 MB, and VCF runs with fixed 64 MB blocks unless a budget is given. The runs reported here used those defaults, except FASTQ, which was given 1000 MB; Supplementary Tables S5 and S6 add VCF under an explicit 2000 MB budget.

### 2.4 Losslessness and benchmarking

Every reported measurement is paired with a full decompression and a byte-level comparison against the original. The tools expose this as a *--verify flag*, and every compressed stream is an OpenZL frame carrying checksums of both its compressed bytes and its decompressed content (OpenZL’s default, which NYX keeps enabled); NYX’s own container headers carry no separate checksum. In a test of 600 random single-bit corruptions, 150 in an archive of each format, every one caused decompression to fail and none produced incorrect output. Beyond the benchmark inputs, a validation corpus exercises empty records, missing trailing newlines, CRLF line endings, ragged line widths, dense IUPAC ambiguity codes and soft-masked repeats. Wall-clock time and peak resident set size are read from /usr/bin/time, and each tool is run once per file.

## 3 Experimental Results

To ensure consistency, all measurements reported were performed on a single TACC Lonestar6 vm-small node (16 cores, 31 GB of RAM), and every multi-threaded tool was given the same 16 threads. The following parameters were used: pigz -9 -p16 [20]; zstd -19 --long=27 -T16 [21]; xz -9e -T16 [22]; 7zz a -t7z -mx=9 -mmt=16 [23]; brotli -q 11 [24] and bgzip -l 9 -@16 [25] for BED; bcftools view -Ob -l9 --threads 16 [26]; spring -c -t 16; and NYX with --threads 16. NAF is single-threaded by design and is reported as such. All formats compress through one OpenZL 0.2.5 build. Genozip [9, 10] is closed-source and licence-gated; we report an open-source comparison set. Τhe comparison can be reproduced by following the steps outlined in the GitHub repo linked in the Data Availability section. GTShark [27] and VCFShark [28] are discussed in section 3.3 but not tabulated, for reasons given there.

Baseline arguments are identical across formats, and NYX compresses against shipped pre-trained models with no training at compression time. Ratio is uncompressed bytes over compressed bytes, while throughput is uncompressed bytes over wall-clock seconds, and peak memory is the peak resident set size (RSS): the most physical memory the process held at any moment, as reported by /usr/bin/time. Decompression includes the post-processing that restores the exact original layout. All decompressed outputs are byte-for-byte identical to the input.

### 3.1 FASTA

#### 3.1.1 Ratio and throughput

On GRCh38.p14 (Table 1; Figure 1a, b) NYX reaches 4.81x at 204 MB/s, and exceeds the best general-purpose codec (xz at 4.52x) on ratio and on compression speed at the same time. NAF at its strongest setting reaches 4.97x, 3.3% more, but takes 1201 s against NYX’s 16.4 s.

**Figure 1.**
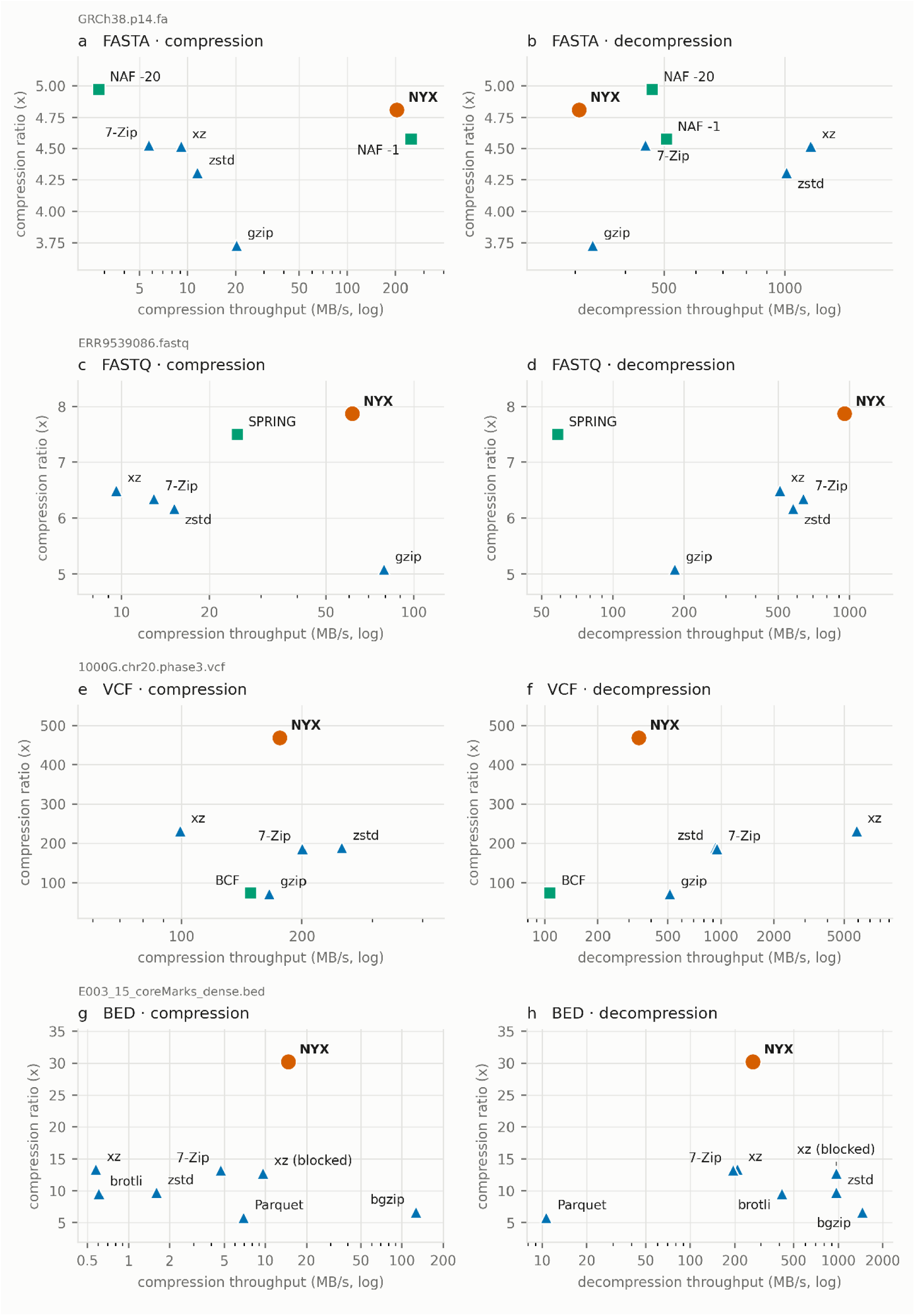
Compression ratio against throughput, one row per format: compression on the left (a, c, e, g), decompression on the right (b, d, f, h), for FASTA (GRCh38.p14), FASTQ (ERR9539086), VCF (1000G chr20) and BED (a ChromHMM chromatin-state segmentation, E003_15_coreMarks_dense). Throughput is log-scaled, so upper-right is better. Colour marks a tool’s family and every point is labelled.

#### 3.1.2 Per-genome results

**Table 1.** Whole-file FASTA results, 16 threads. Ratios for all six codecs.

| Input | GB | NYX | NAF - 20 | NAF - 1 | xz -9e | zstd - 19 | gzip - 9 | NYX s | NAF s | NYX GB |
| --- | --- | --- | --- | --- | --- | --- | --- | --- | --- | --- |
| GRC m39 | 2.77 | 4.823 | 5.004 | 4.489 | 4.517 | 4.314 | 3.580 | 15 | 1036 | 2.6 |
| T2T-CHM1 3v2.0 [29] | 3.16 | 4.430 | 4.660 | 4.331 | 4.203 | 4.032 | 3.382 | 20 | 1170 | 3.0 |
| GRCh 38.p1 4 | 3.35 | 4.811 | 4.971 | 4.576 | 4.518 | 4.309 | 3.729 | 16 | 1201 | 2.6 |
| wheat_IWG SC | 14.79 | 5.491 | 8.278 | 4.512 | 7.200 | 6.151 | 3.608 | 151 | 5408 | 3.9 |

#### 3.1.3 Repetitive assemblies

Bread wheat, with a genome that is roughly 85% repetitive, is the one assembly on which the general-purpose codecs also overtake NYX: NAF -20 reaches 8.28x, xz -9e 7.20x and zstd -19 6.15x, against 5.49x for NYX. The 2-bit packed sequence that NYX produces compresses to 45.9 MB under LZMA but to 85.7 MB under OpenZL’s best LZ configuration, and OpenZL’s untrained, trained and generic profiles all fall within 1% of one another on it. The deficit is therefore a property of the entropy backend rather than of the transform, and would not be recovered by further training. NYX retains its advantage in time and memory on this input, compressing in 151 s and 3.9 GB against 1353 s for xz - 9e and 5408 s for NAF -20.

#### 3.1.4 NAF’s compression level

NAF’s level parameter does not behave monotonically on genomic DNA, which matters for anyone benchmarking it. On a 150 MB slice of GRCm39 the ratio falls from 4.42x at -1 to 4.26x at -5, recovers slowly to 4.38x at -15, then jumps to 4.82x at -20. Level 22 is the highest ratio we measured, but it buys only 0.3% over -20 for 3.9x the memory and 1.2x the time, so we benchmark NAF at -1 and -20.

### 3.2 FASTQ

#### 3.2.1 Ratio and throughput

On ERR9539086 (Table 2; Figure 1c, d) NYX reaches 7.87x against SPRING’s 7.50x, at 62 MB/s against SPRING’s 25 MB/s, and decodes 16.3 times faster.

**Table 2.** Whole-file FASTQ results, 16 threads. . “Best GP” is the highest ratio among gzip -9, zstd - 19 --long=27 and xz -9e for that run. ERR9539086 is the run plotted in Figure 1c, d.

| Input | size (GB) | best GP | SPRING | NYX | SPRING s | NYX s | SPRING GB | NYX GB |
| --- | --- | --- | --- | --- | --- | --- | --- | --- |
| SRR8899104 | 27.97 | 6.837 | 9.627 | 7.278 | 895 | 296 | 5.2 | 2.3 |
| ERR9539079 | 14.64 | 6.957 | 8.975 | 8.769 | 512 | 219 | 8.0 | 2.2 |
| ERR9539093 | 12.02 | 6.824 | 8.288 | 8.298 | 406 | 186 | 7.1 | 2.1 |
| ERR4186945_1 | 11.91 | 7.344 | 12.026 | 8.123 | 483 | 266 | 5.3 | 1.9 |
| ERR9539086 | 8.38 | 6.499 | 7.500 | 7.871 | 336 | 136 | 5.4 | 2.1 |
| DRR206632 | 1.24 | 10.679 | 23.693 | 11.918 | 33 | 28 | 2.1 | 2.1 |

#### 3.2.2 Per-run results

Table 2 reports six Illumina runs. NYX compresses smaller than the best general-purpose codec on every run, and smaller than SPRING on two of the six, all of them libraries without duplicate reads. SPRING’s advantage on the other four is examined in Section 3.2.3.

#### 3.2.3 SPRING results analysis

SPRING and NYX exploit different sources of redundancy in a FASTQ file. SPRING clusters reads that share k-mers, reorders them so that overlapping reads are adjacent, and encodes each read against a consensus of the overlapping reads around it [11]. Its compression ratio therefore increases with the degree to which reads in a library overlap or repeat one another. NYX does not reorder reads: it separates each record into identifier, sequence and quality streams and relies on the sequence coder to find repeats, so it exploits similarity between reads only within that coder’s window.

Supplementary Table S1. reports the fraction of exact-duplicate reads in each library, estimated from the first four million reads, together with the ratio of SPRING’s compression ratio to NYX’s. On the three libraries with no detectable duplicates (ERR9539086, ERR9539079 and ERR9539093) the two tools compress to within 2.3% of one another, NYX smaller on two of them. On the three libraries with duplicate fractions between 4% and 90%, SPRING compressed between 1.3 and 2.0 times smaller. The relationship is not monotonic: the largest advantage was on DRR206632, a small-RNA library with 60% duplicates, rather than on SRR8899104 with 90%. Exact duplication is a lower bound on the redundancy SPRING can exploit, since overlapping but non-identical reads also contribute, and we did not measure the latter directly.

Where SPRING achieved the higher ratio it did so at 1.2 to 3.0 times the compression time and 1.0 to 3.6 times the peak memory (Table 2). For libraries with substantial read duplication, such as small-RNA or amplicon sequencing, SPRING is expected to compress smaller; for libraries without it, NYX matches SPRING’s ratio at a fraction of the time and memory. The duplicate fraction can be estimated from the first few million reads before compression and used to choose between the two.

#### 3.2.4 Memory

Peak resident memory (Table 2; Figure 2) is 1.9-2.3 GB across this entire size range and effectively independent of input size: the 27.97 GB run peaks at 2.3 GB, against 2.1 GB for the 1.24 GB run. SPRING peaks between 2.1 and 8.0 GB on the same runs.

**Figure 2.**
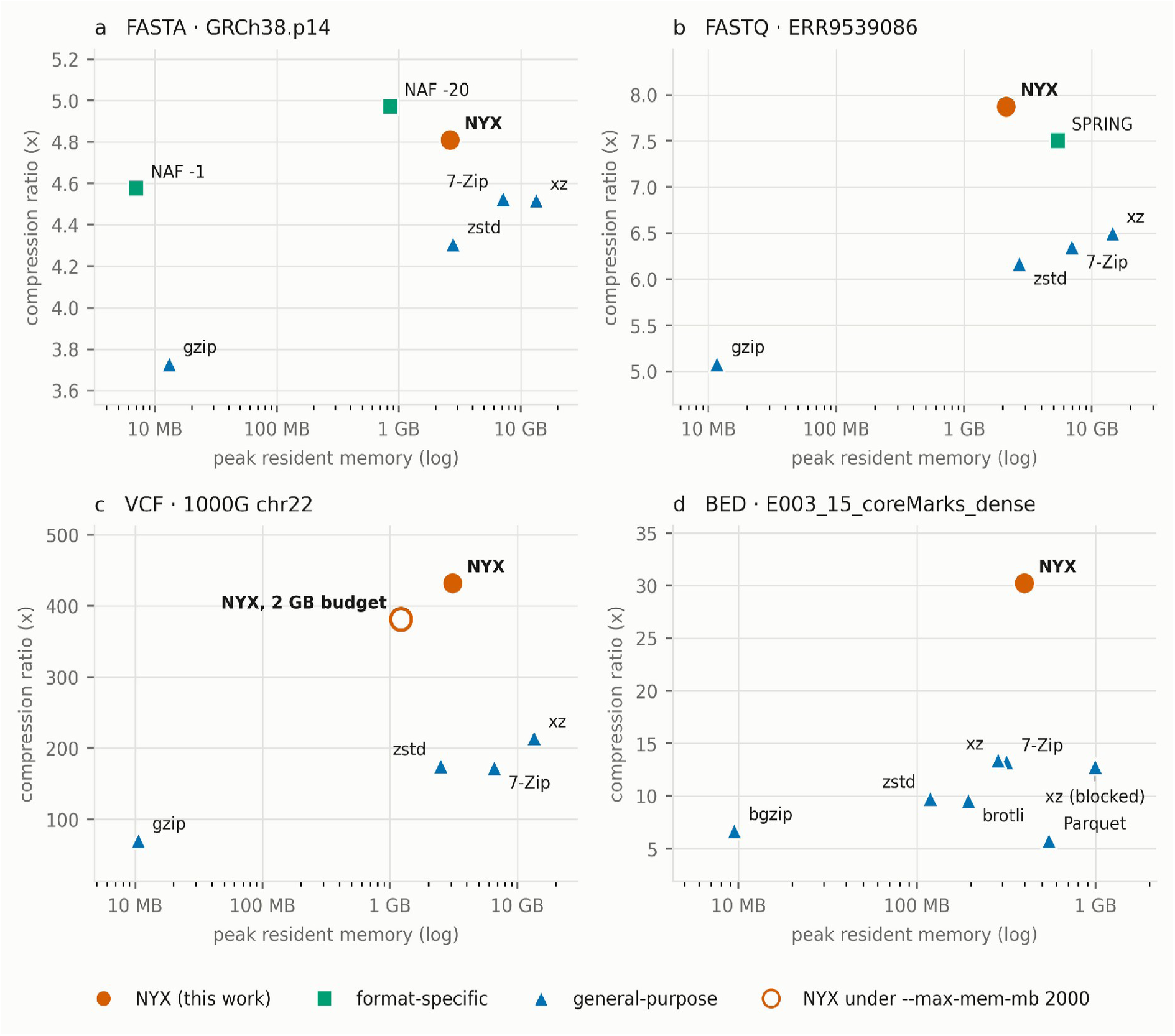
Compression ratio against peak resident memory, one file per format: the Figure 1 file for FASTA, FASTQ and BED, and 1000 Genomes chr22 for VCF, the file on which every tool’s peak memory was measured. Memory is log-scaled, so upper left is better; the hollow marker is NYX under--max-mem-mb 2000. Above and to the left of NYX: NAF -20 on FASTA; no tool in the other panels. Peak memory is measured at 16 threads. For gzip, zstd -19 and xz -9e on FASTA and FASTQ it comes from a separate run with the same invocations, which produced archives byte-identical to those in Tables 1 and 2. The memory budget shown for VCF is --max-mem-mb 2000.

### 3.3 VCF

#### 3.3.1 Ratio

Table 3 reports ten VCFs spanning six structural subtypes, from a 3.6 MB annotation set to an 18.4 GB phased panel. NYX gives the highest ratio on 9 of them. The margin tracks how much of the file is genotype data: on the whole 1000 Genomes phase-3 chromosomes it reaches 468.67x, about 2.02x the ratio of xz -9e, the strongest baseline; on single-sample call sets the margin is about 1.3x; and on a 3.6 MB annotation-only file NYX loses to xz. These are whole chromosomes, not prefixes.

**Table 3.** Whole-file VCF results, 16 threads. Ratio is uncompressed bytes over compressed bytes. The final column is NYX over the best baseline on that file; above 1 means NYX compresses smaller.

| Input | subtype | GB | NYX | gzip | zstd | xz | NYX / best |
| --- | --- | --- | --- | --- | --- | --- | --- |
| 1000G.ch<br>r20.phase<br>3 | panel | 18.42 | 468.67x | 72.62x | 189.45x | 232.32x | 2.02 |
| 1000G.ch<br>r22.phase<br>3 | panel | 11.21 | 431.80x | 70.40x | 175.27x | 214.56x | 2.01 |
| gnomad_<br>v4.1_gen<br>omes_chr<br>Y | sites-<br>frequency | 4.63 | 19.30x | 10.07x | 15.71x | 17.63x | 1.09 |
| HG002_<br>GRCh38_<br>v4.2.1 | single-sample | 2.84 | 49.67x | 22.95x | 34.14x | 38.09x | 1.30 |
| HG004_<br>GRCh38_<br>v4.2.1 | single-sample | 2.78 | 52.15x | 23.38x | 34.67x | 38.71x | 1.35 |
| clinvar_G<br>RCh38 | sites-<br>annotated | 1.94 | 16.86x | 11.00x | 14.46x | 15.36x | 1.10 |
| seqc2_hc<br>c1395_ss<br>nv_super<br>se | somatic | 1.49 | 6.48x | 4.33x | 5.98x | 6.42x | 1.01 |
| seqc2_hc<br>c1395_ss<br>nv | somatic | 0.06 | 30.80x | 14.98x | 24.99x | 27.33x | 1.13 |
| gvcf_nfco<br>re_test2.g | gvcf | 0.04 | 18.32x | 6.88x | 10.70x | 13.16x | 1.39 |
| civic_nigh<br>tly | sites-<br>annotated | 0.00 | 17.18x | 13.69x | 18.35x | 19.75x | 0.87 |

#### 3.3.2 Throughput

On the files above 1 GB, NYX compresses at 77–343 MB/s and decompresses at 341–1112 MB/s. Over the same inputs zstd -19 --long=27 compresses at 11–251 MB/s and xz -9e at 6–99 MB/s. On the five other files above 1 GB, NYX compresses 5.4 to 17 times faster than zstd -19 and 7.5 to 34 times faster than xz -9e. The exception is the two phased 1000 Genomes panels, whose long repeated runs suit zstd’s long-distance matching: there zstd -19 is 1.4 times faster than NYX on chr20 (Figure 1e, f) and level on chr22, at less than half NYX’s ratio, while xz -9e is 1.8 to 2.8 times slower. Absolute throughput falls sharply on small inputs, where fixed start-up costs dominate -- on the 3.6 MB annotation set NYX manages about 10 MB/s against gzip’s hundred -- but the ranking against the strong codecs does not change: on that same file NYX is 5.8 times faster than zstd -19 and 6.7 times faster than xz -9e, because their throughput collapses too. On 1000 Genomes chr22, NYX peaks at 3.11 GB, or 1.22 GB under a 2000 MB budget, against 13.56 GB for xz -9e (Figure 2c; Supplementary Table S6).

#### 3.3.3 Breakdown of transformation contributions

To measure what each component of the VCF transform contributes, we compressed six files representing different VCF subtypes with the transforms enabled cumulatively, in the order in which they are applied (Supplementary Table S2, Figure 3). The baseline is OpenZL’s generic profile applied to the unmodified file. Every configuration was decompressed and verified against the input, so the ablation measures compression ratio only. On the phased 1000 Genomes panel, format awareness as a whole improved the ratio 4.2-fold, from 88.6x to 373.9x. Splitting the record into per-column streams accounted for the first 2.1-fold, and the haplotype PBWT for a further 1.8-fold; the remaining transforms together contributed less than 10%, and the trained models 3%. The PBWT contributed nothing on the other five files. This is expected, since it exploits shared haplotype structure between samples and the other files carry at most two. On the files without a genotype matrix, the gains came from the annotation fields. The per-INFO-key streams were the largest single contributor on ClinVar and on the somatic call set, where INFO carries most of the bytes. The dictionary transform over repeated text was the largest contributor on HG002, improving the ratio 1.4-fold on its own, and on the gVCF the column split (1.5-fold) and then the value transforms (1.4-fold, mostly from delta-coding the monotonically increasing END coordinates) contributed most. The annotation split, which separates VEP- and SnpEff-style records into per-field streams, contributed only on CIViC; on the other files the dictionary transform already captured the repeated annotation values whole, and the split was skipped. Across all six files the trained models contributed between 0% and 3%. The improvement from format awareness is therefore almost entirely due to the transforms rather than to training, which is consistent with the shipped models being applied per stream only where they outperform the generic profile.

**Figure 3.**
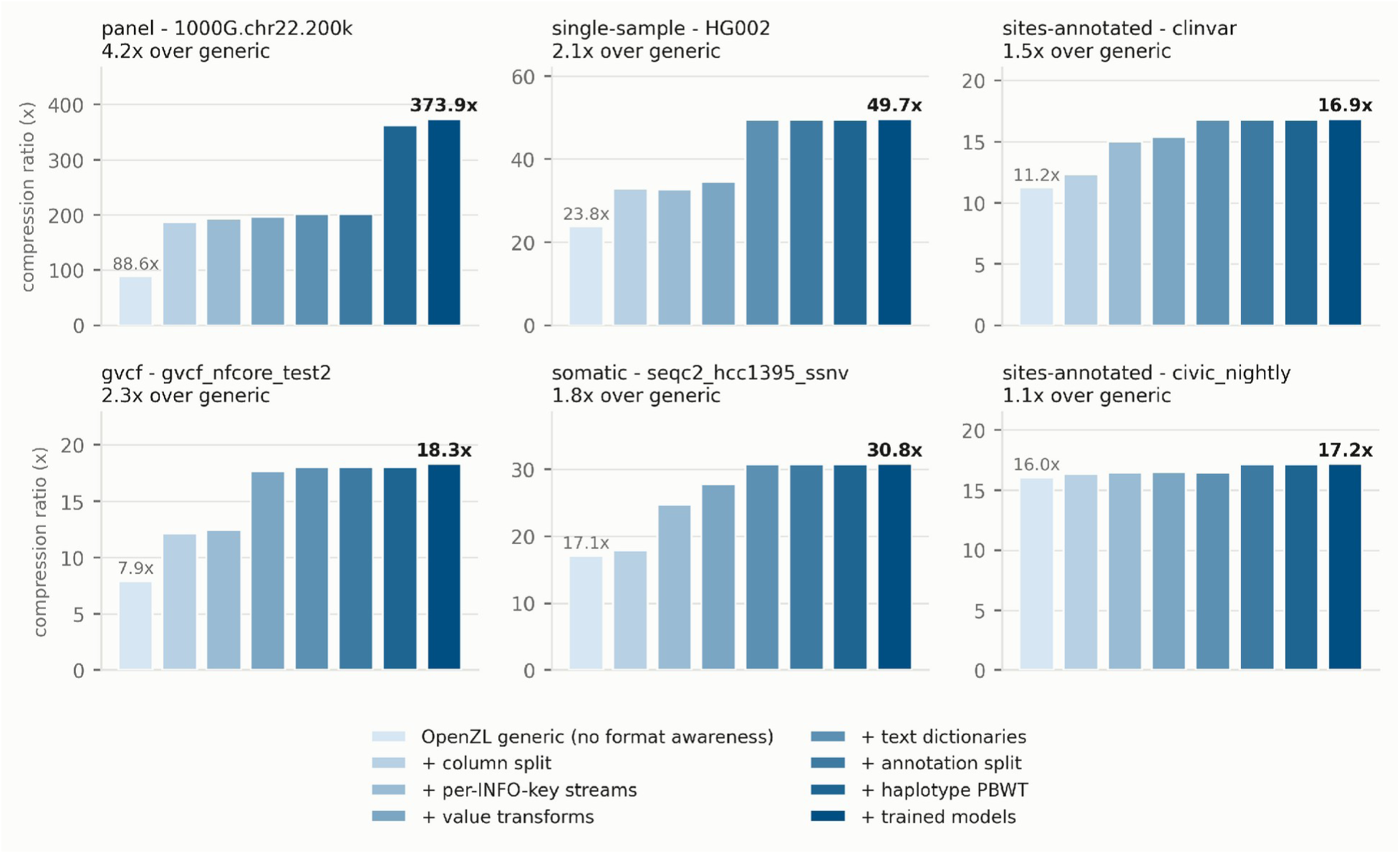
Cumulative contribution of each VCF transform to compression ratio, by file subtype. Bars within each panel add one transform at a time, in the order listed, from OpenZL’s generic profile to the full pipeline. Panel titles give the input file, its subtype and the total gain over the generic profile. The 1000 Genomes panel is the first 200,000 records of chr22; all other inputs are whole files. Y-axis scales differ between panels. Values are in Supplementary Table S2.

#### 3.3.4 GTShark and VCFShark

GTShark [27] and VCFShark [28] are two format-specific VCF compressors that were omitted from this comparison. GTShark is lossless for the genotypes it stores but is not a whole-file compressor: its compress-db mode discards CHROM, REF, ALT, QUAL, FILTER and INFO, so its ratio is not comparable with a lossless compressor. We were not able to successfully run VCFShark 1.1 on our inputs under both gcc 9.4 and 12.2.

### 3.4 BED

#### 3.4.1 Corpus

The BED corpus is 34 public files totalling 1.17 GB, in eight dialects whose widths run from 4 to 15 columns: narrowPeak, broadPeak and gappedPeak peak calls and ChromHMM dense and segment files from the Roadmap Epigenomics consolidated epigenomes [30, 31], the SCREEN registry of candidate cis-regulatory elements [32], the ENCODE transcription-factor binding-site clusters [33] and the FANTOM5 CAGE peak atlas [34]. Roadmap files were taken as every k-th entry of each sorted directory listing, so the selection is reproducible and not picked for results. The widths are not incidental. In a survey of the first 96 KB of 240 public BED-family files we found column counts of 4, 9, 10, 12 and 15 and none at 3, 5, 6, 7 or 8, so the canonical layouts are not the common ones. The format-aware BED path in Genozip [10], for instance, accepts 3, 4, 5, 6, 8, 9 or 12 columns and passes any other width to its generic codec (source code, version 15.0.91); 15 of the 34 files here are narrowPeak or gappedPeak and would not reach it.

#### 3.4.2 Ratio

NYX gives the highest ratio of any tool on all 34 files (Table 4; per-file results in Supplementary Table S3), at 1.24 to 2.30 times the best of zstd -19, xz -9e, 7-Zip and brotli -q 11 on each file (median 1.38). Over the whole corpus that is 9.05-fold against 6.19 for xz -9e, 6.09 for 7-Zip and 4.06 for bgzip, the block-gzip format used for indexed BED files [25]. The gain is largest on ChromHMM dense files (29.5-fold against 13.3; Figure 1g, h), where thickStart, thickEnd and itemRgb are stored as references, and smallest on gappedPeak (4.45 against 3.16), whose block lists and three floating-point statistics leave the least structure between columns. Figure 4a shows the advantage file by file: it is present at every width.

**Figure 4.**
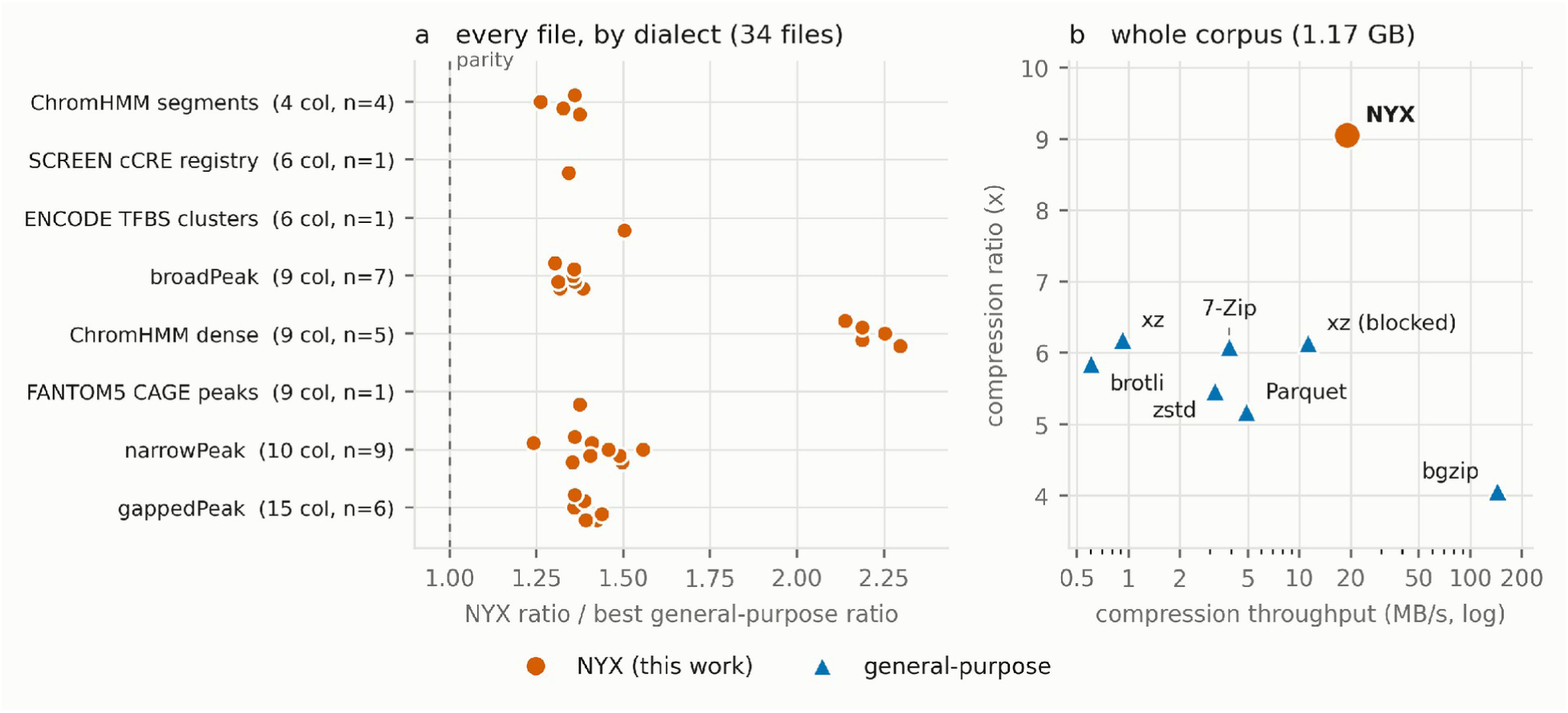
BED across the whole corpus. (a) For each of the 34 files, NYX’s compression ratio divided by that of the best of zstd -19, xz -9e (single-stream or blocked), 7-Zip and brotli -q 11 on the same file, grouped by dialect with its column count; the dashed line is parity, and every point lies to its right. (b) Ratio against compression throughput for the corpus as a whole, one point per tool: total uncompressed over total compressed bytes, and total bytes over total seconds. pigz -9 is omitted because it coincides with bgzip.

**Table 4.**
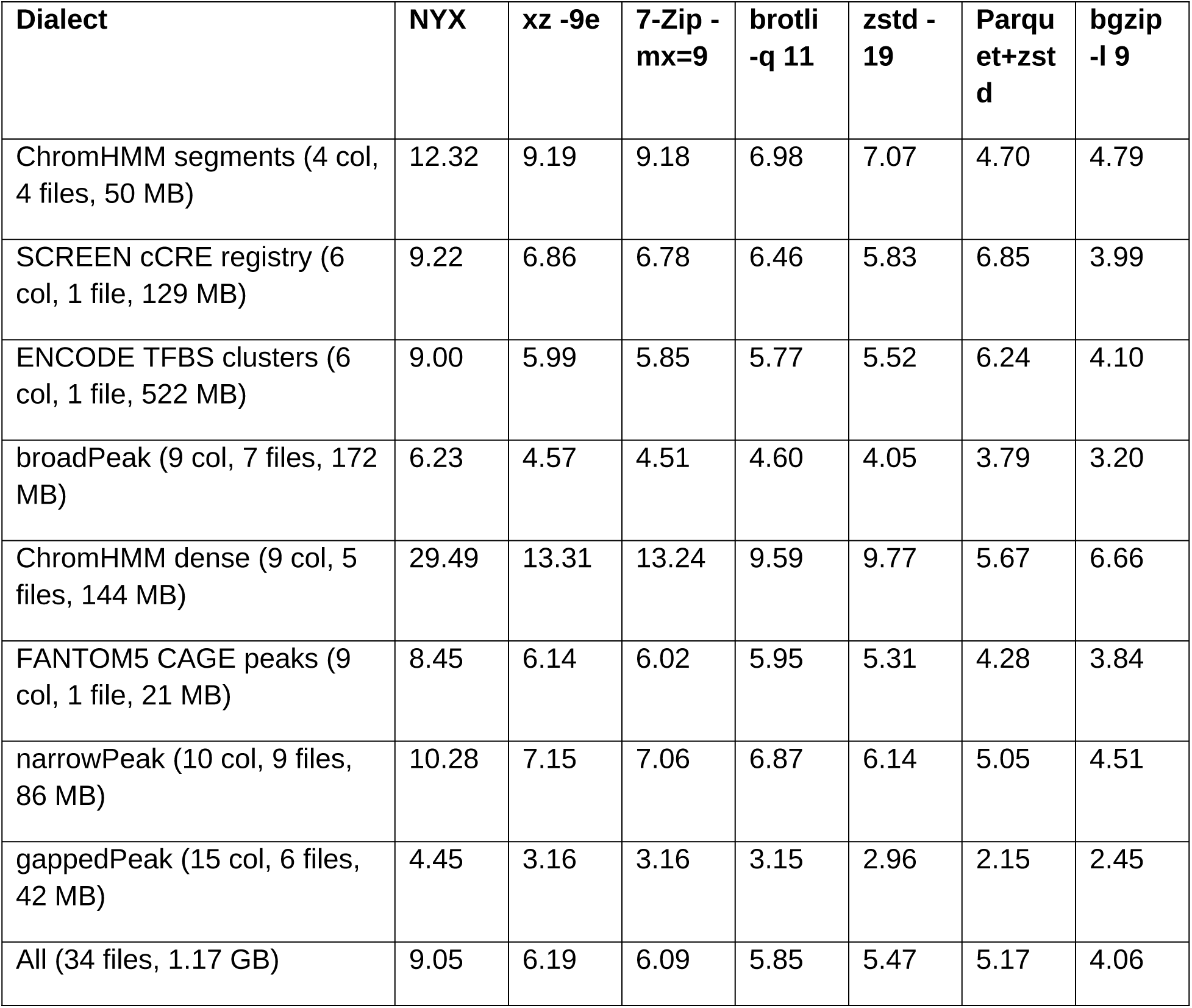
BED compression ratio by dialect, 16 threads, node-local output. Each entry is total uncompressed over total compressed bytes for the files of that dialect, so larger files weigh more; Supplementary Table S3 gives every file, including xz with a blocked stream and pigz -9. Every entry, NYX’s and the baselines’, was decompressed and compared byte-for-byte with its input.

Splitting a table into columns is not by itself new, so we also compressed every file with Apache Parquet [35] at zstd level 19 with dictionary encoding, typing a column as integers wherever that round-trips exactly -- the columnar format an engineer would reach for, held to the same byte-exact standard. NYX is 1.35 to 5.33 times smaller (median 2.04); Parquet’s corpus ratio, 5.17, is below that of xz -9e. What carries the result is the choice of encoding per column and the references between columns, not the split.

#### 3.4.3 Throughput and memory

Speed is a weaker argument for BED than for the other formats, and we state where it fails. NYX compresses faster than zstd -19 on all 34 files, faster than 7-Zip on 24 and faster than single-stream xz -9e on all 34. xz -9e cannot use more than one thread on a file below 192 MiB, which is 33 of the 34 files here, so Supplementary Tables S3 and S4 also report it with its block size set to one sixteenth of the input, which lets it use all 16 threads at a cost of 0.8% of its corpus ratio. That configuration compresses faster than NYX on 27 of the 34 files, all between 3 and 53 MB, where NYX’s per-file calibration is a large share of the work. Over the corpus as a whole NYX is still the faster of the two (61 s against 104 s). NYX is not dominated on any file: no tool is both faster and smaller on any of the 34 (Figure 4b). Decompression is mid-table: NYX decompresses the corpus in 2.6 s. bgzip -l 9 (0.7 s), zstd -19 (1.4 s) and blocked xz -9e (1.6 s) are faster; pigz -9 (3.1 s), brotli -q 11 (3.5 s), xz -9e (8.1 s), 7-Zip -mx=9 (8.9 s) and Parquet+zstd (98.8 s) are slower. Peak memory has a median of 430 MB per file and a maximum of 3.0 GB, against 227 MB and 5.3 GB for blocked xz (Figure 2d).

### 3.5 Overall comparison

Across the 54 inputs, 4 genome assemblies, 6 sequencing runs, 10 VCF files and 34 BED files, NYX compressed smaller than both strong general-purpose codecs, zstd -19 and xz -9e, on 52, and produced the smallest file of any codec tested on 45. It compressed faster than xz -9e on every FASTA, FASTQ and VCF input, and faster than zstd -19 on all of them except the 1000 Genomes chr20 panel, where zstd -19 is 1.4 times faster at less than half NYX’s ratio. On BED, where most files are a few tens of megabytes, xz with a blocked stream compresses faster than NYX on 27 of 34, at a lower ratio (Section 3.4.3). Against the format-specific tools it was within 3-5% of NAF on the mammalian assemblies while compressing 57-73 times faster, and matched SPRING to within 2.3% on the FASTQ libraries without duplicate reads in 2.2 to 2.5 times less time and with up to 3.6 times less memory. No codec tested produced a smaller file than NYX in less time on any of the 54 inputs. NYX was also the only tool in the comparison to handle all four formats, to run against fixed pre-trained models without a training pass on the user’s data, and to bound its peak memory independently of input size.

## 4 Limitations

We would like to acknowledge several limitations. Firstly, the throughput advantage narrows on small inputs. Below roughly 100 MB, fixed start-up costs dominate and the choice of compressor has little practical consequence. Secondly, the included models were trained on a heterogeneous corpus spanning multiple species, sequencing platforms and VCF subtypes, and each stream is compressed with the included model only where it beats OpenZL’s generic profile. This bounds the cost of a poor fit, but does not eliminate it. A file whose statistics lie well outside the training corpus, will compress with a ratio below what a model trained on similar data would reach. The training scripts are included in the distribution for users who wish to close that gap on their own data.

## 5 Conclusion

NYX compresses FASTA, FASTQ, BED and VCF files through a single OpenZL substrate, with a reversible transform per format and a fixed set of pre-trained models, so that no training is performed on the user’s data. On the 54 inputs benchmarked here it exceeds the strongest general-purpose codecs on compression ratio on 52 and on both ratio and speed on the FASTA, FASTQ and VCF inputs, and no codec tested produces a smaller file in less time. Every reported result was decompressed and compared byte-for-byte against its source, and inputs the encoder cannot model are stored verbatim, so the round trip is exact regardless of content. The source code, models, benchmark harness and all scripts needed to regenerate the tables and figures are publicly available.

## Data availability

All inputs are public. The VCF, FASTA and FASTQ accessions and download scripts, the shipped models, the benchmark harness, the round-trip test corpora and the scripts that regenerate every figure and table in this manuscript can be found in https://github.com/Georgakopoulos-Soares-lab/compression.

## Supplementary Tables

Byte-exact, format-aware compression for FASTA, FASTQ, BED and VCF files

**Supplementary Table S1.** Exact-duplicate read fraction against the SPRING ratio advantage, measured on the first four million reads of each library (scripts/fastq/duplication_profile.sh).

| run | duplicate reads | NYX | SPRING | SPRING / NYX |
| --- | --- | --- | --- | --- |
| SRR8899104 | 89.6% | 7.28x | 9.63x | 1.32x |
| DRR206632 | 59.9% | 11.92x | 23.69x | 1.99x |
| ERR4186945_1 | 4.2% | 8.12x | 12.03x | 1.48x |
| ERR9539086 | 0.0% | 7.87x | 7.50x | 0.95x |
| ERR9539079 | 0.0% | 8.77x | 8.97x | 1.02x |
| ERR9539093 | 0.0% | 8.30x | 8.29x | 1.00x |

**Supplementary Table S2.** Cumulative contribution of each VCF transform, by subtype. Each row adds one transform to the row above.

| transform<br>(cumulative) | 1000G.chr22.2<br>00k | HG002 | clinvar | gvcf | seqc2_hcc139<br>5_ssnv | civic |
| --- | --- | --- | --- | --- | --- | --- |
| OpenZL generic<br>(no format<br>awareness) | 88.59x | 23.79x | 11.23x | 7.91x | 17.09x | 16.04x |
| + column split | 187.29x | 32.92x | 12.34x | 12.11x | 17.89x | 16.32x |
| + per-INFO-key<br>streams | 192.94x | 32.74x | 15.04x | 12.46x | 24.74x | 16.46x |
| + value<br>transforms | 196.36x | 34.62x | 15.39x | 17.67x | 27.82x | 16.50x |
| + text<br>dictionaries | 201.44x | 49.51x | 16.82x | 18.05x | 30.75x | 16.47x |
| + annotation<br>split | 201.44x | 49.51x | 16.82x | 18.05x | 30.75x | 17.16x |
| + haplotype<br>PBWT | 362.45x | 49.51x | 16.82x | 18.05x | 30.75x | 17.16x |
| + trained<br>models | 373.94x | 49.67x | 16.86x | 18.32x | 30.80x | 17.18x |

**Supplementary Table S3.**
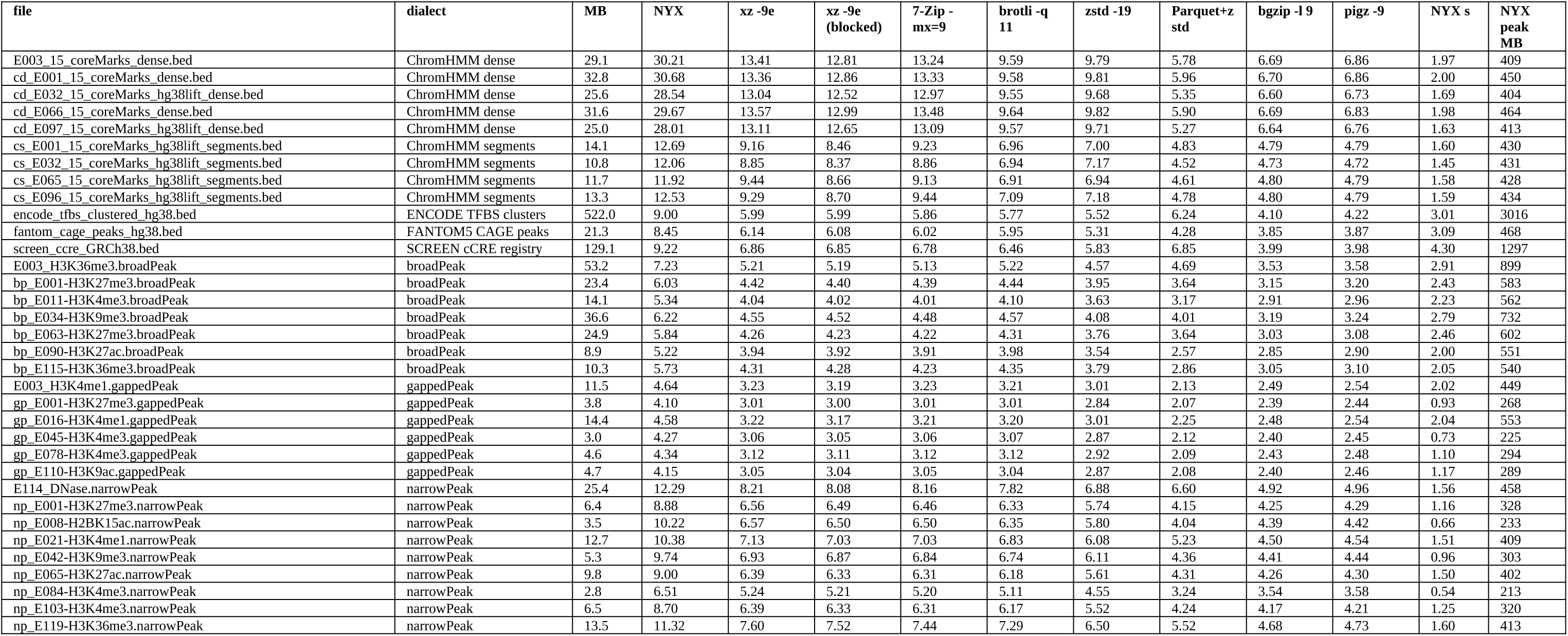
Per-file BED results, 34 files, 16 threads, one TACC Lonestar6 vm-small node (16 cores, 31 GB). Ratio is uncompressed over compressed bytes; NYX time and peak are for compression. Every entry was decompressed and compared byte-for-byte with its input. Source: results/bed_bench.csv in the repository.

**Supplementary Table S4.**
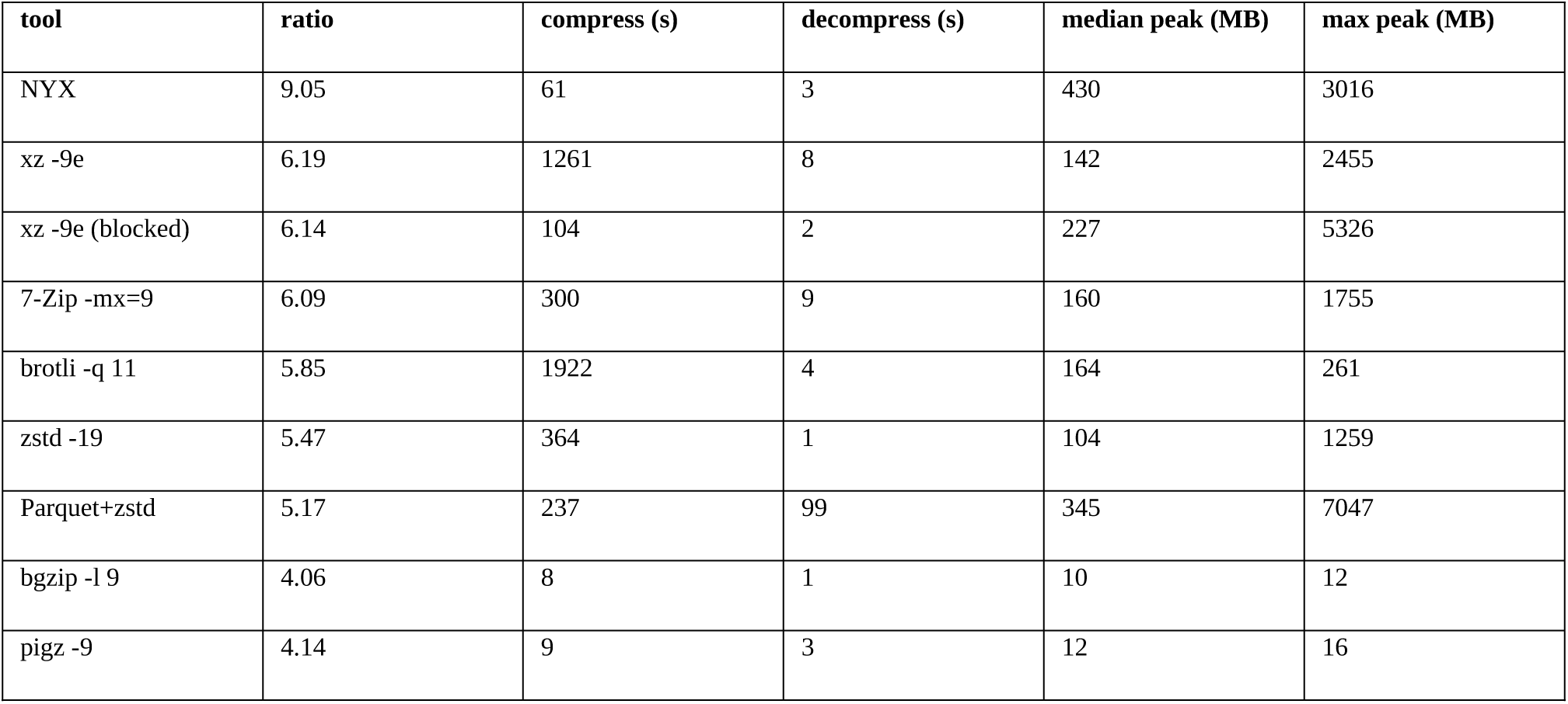
The BED corpus as a whole (34 files, 1.17 GB), 16 threads. Times are sums over files; peak resident memory is per file. Archives and round-trip outputs were written to node-local disk for every tool. xz -9e (blocked) is xz -9e -T16 with --block-size set to input size / 16.

**Supplementary Table S5.**
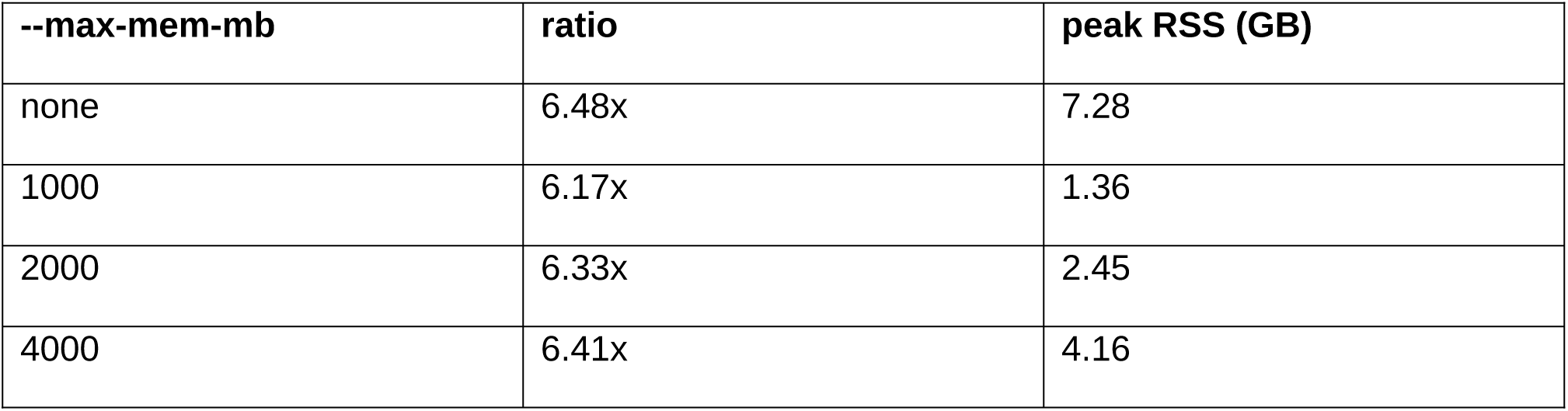
Compression ratio and measured peak RSS against the requested memory budget (seqc2_hcc1395_ssnv_superset.vcf, 1.49 GB, 16 threads). Every run verified byte-exact.

| --max-mem-mb | ratio | peak RSS (GB) |
| --- | --- | --- |
| none | 6.48x | 7.28 |
| 1000 | 6.17x | 1.36 |
| 2000 | 6.33x | 2.45 |
| 4000 | 6.41x | 4.16 |

**Supplementary Table S6.** Peak resident memory (GB) during compression, 16 threads, same node and inputs as Table 3.

| input | NYX | NYX (2 GB budget) | gzip | zstd -19 | xz -9e |
| --- | --- | --- | --- | --- | --- |
| 1000G.chr22.phase3.vcf | 3.11 | 1.22 | 0.01 | 2.51 | 13.56 |
| clinvar_GRCh38.vcf | 4.93 | 1.91 | 0.01 | 2.60 | 8.29 |
| HG002_GRC h38_v4.2.1.vcf | 5.19 | 1.66 | 0.01 | 2.52 | 11.30 |

